# DeepMinimizer: A Differentiable Framework for Optimizing Sequence-Specific Minimizer Schemes

**DOI:** 10.1101/2022.02.17.480870

**Authors:** Minh Hoang, Hongyu Zheng, Carl Kingsford

## Abstract

Minimizers are k-mer sampling schemes designed to generate sketches for large sequences that preserve sufficiently long matches between sequences. Despite their widespread application, learning an effective minimizer scheme with optimal sketch size is still an open question. Most work in this direction focuses on designing schemes that work well on expectation over random sequences, which have limited applicability to many practical tools. On the other hand, several methods have been proposed to construct minimizer schemes for a specific target sequence. These methods, however, require greedy approximations to solve an intractable discrete optimization problem on the permutation space of *k*-mer orderings. To address this challenge, we propose: (a) a reformulation of the combinatorial solution space using a deep neural network re-parameterization; and (b) a fully differentiable approximation of the discrete objective. We demonstrate that our framework, DeepMinimizer, discovers minimizer schemes that significantly outperform state-of-the-art constructions on genomic sequences.

## 1 Introduction

Minimizers [15, 16] are deterministic methods to sample *k*-mers from a sequence at approximately regular intervals such that sufficient information about the identity of the sequence is preserved. Sequence sketching with minimizers is widely used to reduce memory consumption and processing time in bioinformatics programs such as read mappers [7, 10], *k*-mer counters [3, 5] and genome assemblers [17]. Given a choice of *k*-mer length *k* and window length *w*, a minimizer selects the lowest priority *k*-mer from every overlapping window in the target sequence according to some total ordering *π* over all *k*-mers. Minimizer performance is measured by its density [12] on a target sequence, which is proportional to the induced sketch size.

Depending on the choice of *π*, the resulting density can significantly vary. The theoretical lower-bound of density achievable by any minimizer scheme is given by 𝒪 (1*/w*) [12]. On the other hand, a random initialization of *π* will yield an expected density of 𝒪 (2*/w*) [16], which is frequently used as a baseline for comparing minimizer performance. This motivates the question: How do we effectively optimize *π* to improve the performance of minimizers?

An exhaustive search over the combinatorial space of *π* suffices for very small *k*, but quickly becomes intractable for values of *k* used in practice (i.e., *k ≥* 7) (Section 3.1). To work around this, many existing approaches focus on constructing minimizer schemes from mathematical objects with appealing properties such as universal hitting sets (UHS) [11, 4, 12, 14, 19]. While these schemes provide upper-bound guarantees for expected densities on random sequences, they only obtain modest improvements over a random minimizer when used to sketch a specific sequence [19].

The idea of learning minimizer schemes tailored towards a target sequence has been previously explored, although to a lesser extent. Current approaches include heuristic designs [1, 8], greedy pruning [2] and construction of *k*-mer sets that are well-spread on the target sequence [20]. However, these methods only learn crude approximations of *π* by dividing *k*-mers into disjoint subsets with different priorities to be selected. Within each subset, the relative ordering is arbitrarily assigned to recover a valid minimizer, hence they are not necessarily optimal. We give a detailed overview of these methods in Section 2.

This paper instead tackles the problem of directly learning a total order *π*. The hardness of solving such a task comes from two factors, which we will review in detail in Section 3.1: (1) the search space of *k*-mer orderings is very large; and (2) the density minimizing objective is discrete. To overcome the above challenges, we propose to reformulate the original problem as parameter optimization of a deep learning system. This results in the first fully-differentiable minimizer selection framework that can be efficiently optimized using gradient-based learning techniques. Our contributions are:

– We define a more well-behaved search space that is suitable for gradient-based optimization. This is achieved by implicitly representing *k*-mer orderings as continuous score assignments. The space of these assignments is parameterized by a neural network called PriorityNet, whose architecture guarantees that every output assignment is *consistent* (i.e., corresponding to valid minimizer schemes). The modelling capacity of PriorityNet can be controlled via increasing its architecture depth, which implies a mild restriction on the candidate space in practice (Section 3.2).
– We approximate the discrete learning objective by a pair of simpler tasks. First, we design a complementary neural network called TemplateNet, which outputs potentially *inconsistent* assignments (i.e., template) with guaranteed low densities on the target sequence (Section 3.4). We then search for *consistent* assignments (i.e., valid minimizers) around these templates, which potentially will yield similar densities. This is achieved via a fully differentiable proxy objective (Section 3.3) that minimizes a novel divergence (Section 3.5) between these networks.
– We compare our framework, DeepMinimizer, against various state-of-the-art benchmarks and observe that DeepMinimizer yields sketches with significantly lower densities on genomic sequences (Section 4).

## 2 Related Work

### UHS-based methods

Most existing minimizer selection schemes with performance guarantees over random sequences are based on the theory of universal hitting sets (UHS) [11, 14]. Particularly, a (*w, k*)-UHS is defined as a set of *k*-mers such that every window of length *w* (from any possible sequence) contains at least one of its elements. Every UHS subsequently defines a family of corresponding minimizer schemes whose expected densities on random sequences can be upper-bounded in terms of the UHS size [12]. As such, to obtain minimizers with provably low density, it suffices to construct small UHS, which is often the common learning objective of many existing approaches [12, 4, 19].

In the context of *sequence-specific* minimizers, there are several concerns with this approach. First, the requirement of UHS to “hit” all windows of *every possible* sequence is often too strong with respect to the need of sketching a specific string and results in sub-optimal universal hitting sets [20]. Additionally, since real sequences rarely follow a uniform distribution [18], there tends to be little correspondence between the provable upper-bound on expected density and the actual density measured on a target sequence. In practice, the latter is usually more pessimistic on sequences of interest, such as the human reference genome [19, 20], which drives the development of various *sequence-specific* minimizer selection methods.

### Heuristic methods

Several minimizer construction schemes rank *k*-mers based on their frequencies in the target sequence [1, 8], such that rare *k*-mers are more likely to be chosen as minimizers. These constructions nonetheless rely on the assumption that rare *k*-mers are spread apart and ideally correspond to a sparse sampling. Another greedy approach is to sequentially remove k-mers from an arbitrarily constructed UHS, as long as the resulting set still hits every *w*-long window on the target sequence [2]. Though this helps to fine-tune a given UHS with respect to the sequence of interest, there is no guarantee that such an initial set will yield the optimal solution after pruning.

### Polar set construction

Recently, a novel class of minimizer constructions was proposed based on polar sets of *k*-mers, whose elements are sufficiently far apart on the target sequence [20]. The sketch size induced by such a polar set is shown to be tightly bounded with respect to its cardinality. This reveals an alternate route to low-density minimizer schemes through searching for the minimal polar set. Unfortunately, this proxy objective is NP-hard and currently approximated by a greedy construction [20].

### Remark

In all of the above methods, the common objective to be optimized can be seen as a partition of the set of all *k*-mers into disjoint subsets. For example, frequency values are used to denote different buckets of *k*-mers [1, 8]. Others [19, 2, 4, 20] employ a more fine-grained partitioning scheme defined by the constructed UHS/polar set. Each subset has an assigned priority value, such that *k*-mers from higher priority subsets are always chosen over *k*-mers from lower priority subsets. However, it remains inconclusive how *k*-mers from within the same subset can be optimally selected to recover a total ordering *π*. Practically, these methods resort to using a pre-determined arbitrary ordering to resolve such situations. In contrast, our work investigates a novel approach to directly learn this ordering.

## 3 Methods

### 3.1 Background

Let *Σ* be an alphabet of size |*Σ*| = *σ* and *S* be a sequence containing exactly *l* overlapping *k*-mers defined on this alphabet, i.e., *S ∈ Σ*^*l*+*k*−1^. For some *w* ∈ ℕ^+^ such that *l ≥ w*, we define a (*w, k*)-window as a substring in *S* of length *w* + *k* − 1, which contains exactly *w* overlapping *k*-mers. For ease of notation, we further let *l*_*w*_ ≜ *l* − *w* + 1 denote the number of (*w, k*)-windows in *S*. For the rest of this paper, we assume that *w* and *k* are fixed and given as application-specific parameters.

#### Definition 1. (Minimizer)

A minimizer scheme *m* : *Σ*^*w*+*k*−1^ *→* [1, *w*] is uniquely specified by a total ordering *π* on *Σ*^*k*^. Here, we encode *π* as a function *ρ* : *Σ*^*k*^ *→* ℕ^+^ that maps *k*-mers to its position in *π*. Given a (*w, k*)-window *ω, m* then returns the smallest *k*-mer in *ω* according to *ρ*:

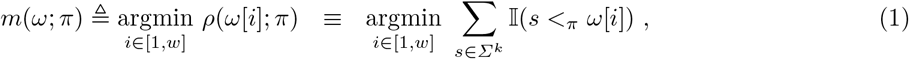

where 𝕀 denotes the indicator function, *ω*[*i*] denotes the *i*-th *k*-mer in *ω*, and *s <*_*π*_ *ω*[*i*] implies *s* precedes *ω*[*i*] in *π*. We break ties by prioritizing *k*-mers that occur earlier in (i.e., to the left of) the window.

When applied to a sequence *S*, the above scheme selects one *k*-mer position from every overlapping window to construct the sequence sketch ℒ (*S*; *m*) = {*t* + *m*(*ω*_*t*_) | *t* ∈ [1, *l*_*w*_]}, with *ω*_*t*_ denoting the *t*^th^ window in *S*. Naturally, a smaller sketch leads to more space and cost savings. As such, we measure minimizer performance by the density factor metric 𝒟 (*S*; *m*) ≜ |ℒ (*S*; *m*) | × (*w* + 1)*/l*_*w*_, which approximates the number of *k*-mers selected per window [12]. The minimizer selection problem is then formalized as density minimization with respect to *π*:

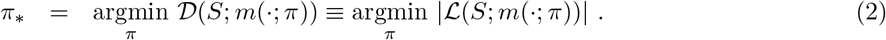

This objective, however, is intractable to optimize for two reasons. First, the number of all *k*-mer permutations scales super-exponentially with *k* and *σ* (i.e., *σ*^*k*^!), thus renders any form of exhaustive search on this space impossible under most practical settings. Furthermore, the set counting operation | ℒ (*S*; *m*(·; *π*))| is non-differentiable even if the solution space is continuous, which makes efficient gradient-based optimizers inaccessible. The remainder of this section therefore proposes a deep-learning strategy to address both these challenges, and is organized as follows.

Section 3.2 describes a unifying view of existing methods as reparameterizations of *ρ* (Definition 1). We then propose a novel deep parameterization called PriorityNet, which relaxes the permutation search space of Eq. 2 into a well-behaved weight space of a neural network.

Section 3.3 shows that density optimization with respect to PriorityNet can be approximated by two sub-tasks via introducing another complementary network, called TemplateNet. This approximation can be formalized as a fully-differentiable proxy objective that minimizes divergence between TemplateNet and PriorityNet.

Section 3.4 and Section 3.5 then respectively discuss the parameterization of TemplateNet and the divergence measure in our proxy objective, thus completing the specification of our framework, DeepMinimizer. An overview of our framework is given in Fig. 1.

**Fig. 1.**
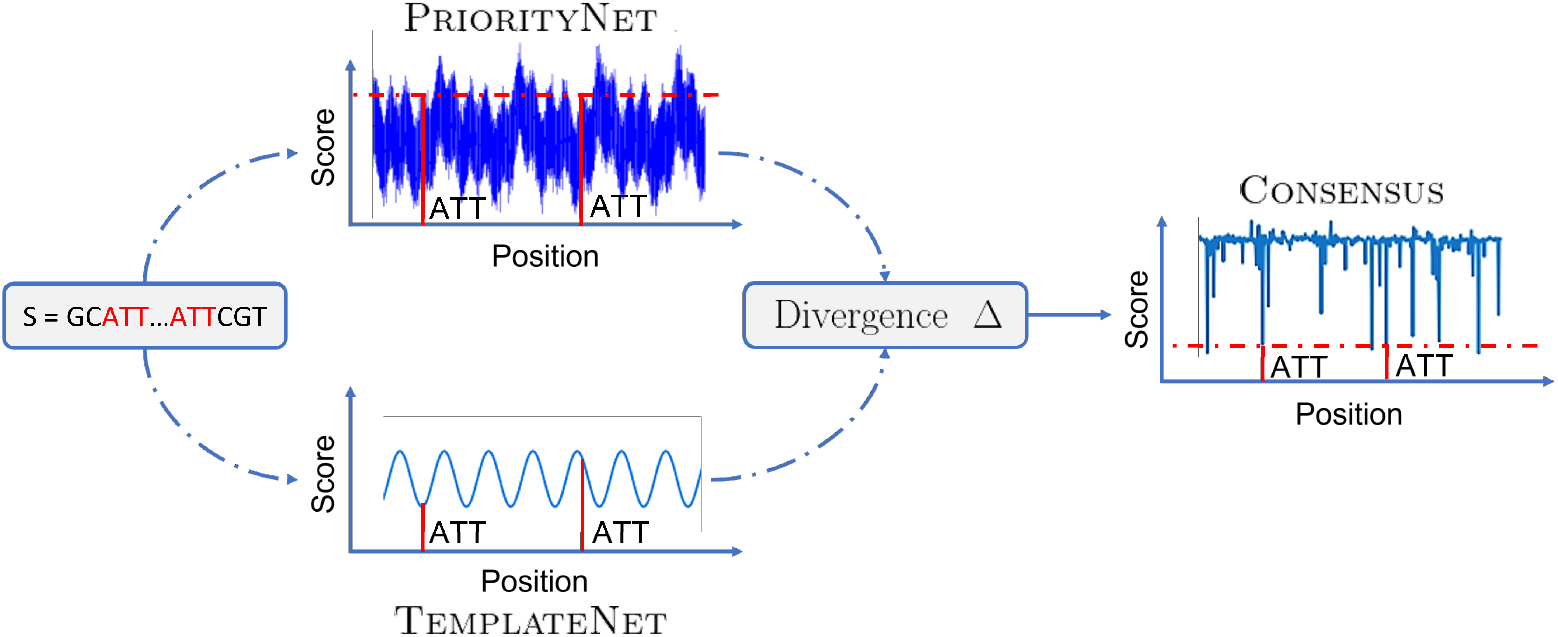
Our DeepMinimizer framework employs a twin network architecture. P riorityNet generates valid minimizers, but has no guarantee on density. In contrast, TemplateNet generates low-density templates that might not correspond to valid minimizers. We minimize the divergence between these networks to arrive at consensus minimizers with low densities on the target sequence.

### 3.2 Search Space Reparameterization

We first remark that many existing methods can be seen as different re-parameterizations of *ρ* in Definition 1. For example, *ρ* can be parameterized with frequency information from the target sequence [1, 8], i.e., 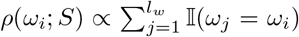; or instantiated with a UHS *υ* [4, 19], i.e., *ρ*(*ω*_*i*_; *υ*) = 𝕀(*ω*_*i*_ *∉ υ*). Similar set-ups have been explored in the context of sequence-specific minimizers using a pruned UHS *υ*(*S*) [2] and a polar set *ζ*(*S*) [20] constructed for the target sequence. We note that there are fewer discrete values potentially assigned by *ρ* than the total number of *k*-mers in all these re-parameterizations. As such, these methods still rely on a pre-determined arbitrary ordering to break ties in windows with two or more similarly scored *k*-mers. When collisions occur frequently, this could have unexpected impact on the final density.

DeepMinimizer instead employs a continuous parameterization of *ρ* using a feed-forward neural network parameterized by weights *α*, which takes as input the multi-hot encoding of a *k*-mer (i.e., a concatenation of its character one-hot encodings) and returns a real-valued score in [0, 1]. This continuous scheme practically eliminates the chance for scoring collisions. Furthermore, the solution space of this re-parameterization is only restricted by the modelling capacity encoded by our architecture weight space. This limitation quickly diminishes as we employ sufficiently large number of hidden layers in the network. We can subsequently rewrite Eq. 2 as optimizing a neural network with density as its loss function:

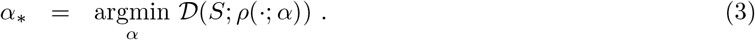

Applying this network on every *k*-mer along *S* can be compactly written as a convolutional neural network, denoted by *f*, which maps the entire sequence *S* to a *score assignment* vector. We require this score assignment to be *consistent* across different windows in order to recover a valid ordering *π* from such implicitly encoded *ρ*. Specifically, one *k*-mer can not be assigned different scores at different locations in *S*. To enforce this, we let the first convolution layer of our architecture, PriorityNet, have kernel size *k*, and all subsequent layers to have kernel size 1. An illustration for PriorityNet when *k* = 2 is given in Fig. 2.

**Fig. 2.**
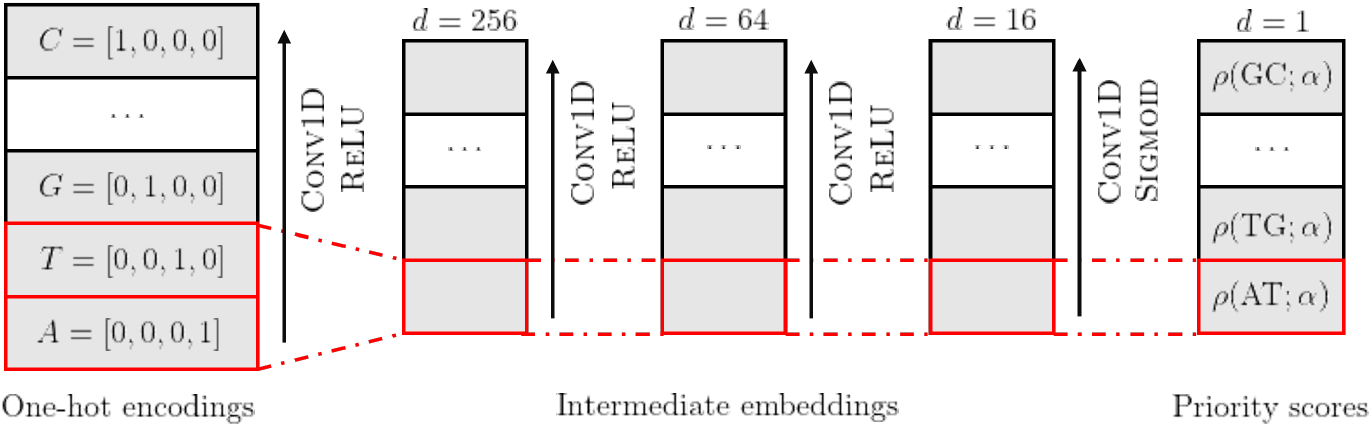
Our PriorityNet architecture for *k* = 2, parameterized by weights *α*, maps sequence multi-hot encoding to priority scores through a series of 3 convolution layers with kernel size [*k*, 1, 1] and [256, 64, 16] embedding channels respectively. Fixing network weights *α*, the computation of assigned priority score to any *k*-mer is deterministic given its character one-hot encodings.

### 3.3 Proxy Objective

The density computation in Eq. 3, however, is not differentiable with respect to the network weights. As such, *α* cannot be readily optimized with established gradient back-propagation techniques used in most deep learning methods. To work around this, we introduce a proxy optimization objective that approximates Eq. 3 via coupling PriorityNet with another function called TemplateNet. Unlike the former, TemplateNet relaxes the *consistency* requirement and generates *template* score assignments that might not correspond to valid minimizer schemes. In exchange, such *templates* are guaranteed to yield low densities by design.

Intuitively, the goals of these networks are complementary: PriorityNet generates valid minimizer schemes in the form of *consistent* priority score assignments, whereas TemplateNet pinpoints neighborhoods of low-density score assignments situated around its output templates. This reveals an alternative optimization route where these networks negotiate towards a consensus solution that (a) satisfies the constraint enforced by PriorityNet; and (b) resembles a template in the output space of TemplateNet, thus potentially yielding low density. Let *f* and *g* denote our proposed PriorityNet and TemplateNet, respectively parameterized by weights *α* and *β*, we formalize this objective as minimizing some divergence measure *Δ* between their outputs:

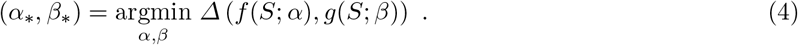

In the remainder of this paper, we detail the full specification of our proxy objective, which requires two other ingredients. First, Section 3.4 discusses the parameterization of our TemplateNet *g* to consistently generate templates that achieve the theoretical lower-bound density [12] on the target sequence. Furthermore, we note that the proxy objective in Eq. 4 will perform best when the divergence measure *Δ* reflects the difference in densities of two score assignments. Section 3.5 then discusses a practical choice of *Δ* to accurately capture high-performing neighborhoods of minimizers. These specifications have strong implications on the expressiveness of the solution space and directly influences the performance of our framework, as shown in Section 4.

### 3.4 Specification of TemplateNet

The well-known theoretical lower bound 1 + 1*/w* for density factor [12] implies that the optimal minimizer, if it exists, samples *k*-mers exactly *w* positions apart. As a result, we want to guarantee that the output of TemplateNet approximates this scenario given any weights initialization. Without loss of generality, we impose that TemplateNet is given by a continuous function *g* : ℝ *→* [0, 1], such that its output template **v** = [*g*(*i*)]_*i*∈[*l*]_ consists of evaluations of *g* restricted to integer inputs (i.e., *k*-mer positions). Then, Proposition 1 below shows a sufficient construction for *g* that approximately yields the optimal density.

#### Proposition 1.

*Let g* : ℝ *→* [0, 1] *be a periodic function with minimal period w, such that g has a unique minimum value on every w-long interval. Formally, g satisfies:*

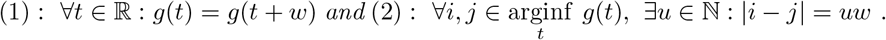

*Then, the template generated by g induces a sketch with density factor* 1 + 1*/w* + *o*(1) *on S when S is sufficiently long (i*.*e*., *l*_*w*_ *≫ w*^2^).

*Proof*. We give a detailed proof of Proposition 1 in Appendix A. □

Note that the resulting sketch induced by *g* does not necessarily correspond to a valid minimizer. While this sketch has low density, it does not preserve the sequence identity like a minimizer sketch, hence is not useful for downstream applications. However, it is sufficient as a guiding template to help PriorityNet navigating the space of orderings.

Proposition 1 leaves us with infinitely many candidate functions to choose from. In fact, TemplateNet can be as simple as *g*(*t*) = sin(2*πt/w*) to generate a near-optimal score assignment. This naïve specification, however, encodes exactly one template (i.e., one that picks *k*-mers from the set of interval positions {*w*, 2*w*, …}), whose proximal neighborhood might not contain any valid minimizertscheme. For example, consider a sequence *S* in which some particular *k*-mer occurs exactly at positions 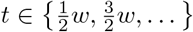. Ideally, we would want to align the *template minima* with these locations, which is not possible given the above choice of *g*. As such, it is necessary that the specification of TemplateNet is sufficiently expressive for Eq. 4 to find an optimal solution.

In particular, we want to construct a parameterized function such that every *k*-mer position can be sampled by at least one sketch encoded in its parameter space. Furthermore, we note that the periodic property is only a sufficient condition to obtain low-density sketches. In practice, we only want the template minima to periodically occur at fixed intervals. Enforcing the scores assigned at all positions to exactly follow a sinusoidal pattern is restrictive and might lead to overlooking good templates. To address these design goals, we propose the following ensemble parameterization:

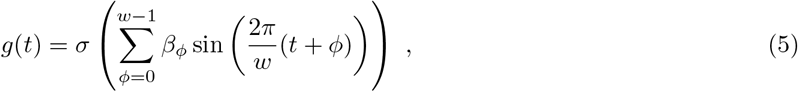

where *σ* denotes a sigmoid activation function, which ensures that *g*(*t*) appropriately maps to [0, 1]; 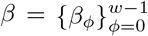 are optimizable parameters such that *β*_*ϕ*_ *≥* 0 and 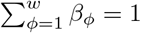.

Optimizing *β* has two implications. First, by adjusting the dominant phase shift *ϕ*_max_ = argmax_*ϕ*_ *β*_*ϕ*_, we can control the offset of the periodic template minima, which leads to good coverage on the target sequence. Second, by adjusting the magnitudes of the remaining phase shifts 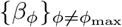, we can have more degrees of freedom to assign scores outside the template minima. Lastly, the non-negative and sum-to-one constraints help to avoid the trivial assignment of squashing all magnitudes to 0 and are easily guaranteed by letting *β* be the output of a softmax layer.

### 3.5 Specification of the Divergence Measure *Δ*

As standard practice, we first consider instantiating *Δ* with the squared *𝓁*^2^-distance. Specifically, let **v**_*f*_ = *f* (*S*; *α*) and **v**_*g*_ = *g*(*S*; *β*) denote the score assignments respectively output by PriorityNet and TemplateNet given *S*, then 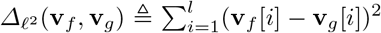. This divergence measure, however, places an excessively strict matching objective at all locations along **v**_*f*_ and **v**_*g*_. Such a perfect match is unnecessary as long as the *k*-mers outside sampled locations are assigned higher scores, and will take away the degrees of freedom needed for the proxy objective to satisfy the constraints implied by PriorityNet.

Consequently, we are interested in constructing a divergence that: (a) strategically prioritizes matching **v**_*f*_ to the minima of the template **v**_*g*_; and (b) enables flexible assignment at other positions to admit more solutions that meet the consistency requirement. To accomplish these design goals, we propose the following asymmetrical divergence:

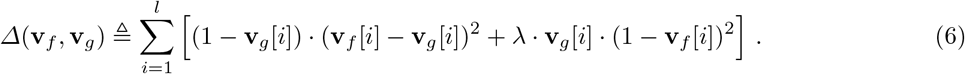

Specifically, the idea behind the first component (1−**v**_*g*_[*i*])·(**v**_*f*_ [*i*]−**v**_*g*_[*i*])^2^ in the summation is to weight each position-wise matching term (**v**_*f*_ [*i*] − **v**_*g*_[*i*])^2^ by its corresponding template score: the weight term (1 − **v**_*g*_[*i*]) implies stronger matching preference around the minima of **v**_*g*_ where the template scores **v**_*g*_[*i*] are low, and vice versa weaker preference at other locations. Furthermore, to ensure that *f* properly assigns higher scores to the locations outside the minima of **v**_*g*_, the second component **v**_*g*_[*i*] · (1 − **v**_*f*_ [*i*])^2^ subsequently encourages *f* to maximize its assigned scores wherever possible, again weighted by the relative relevance of each location. The trade-off between these components is controlled by the hyper-parameter *λ*. Finally, we confirm that this divergence measure is fully differentiable with respect to *α* and *β*, hence can be efficiently optimized using gradient-based techniques. Particularly, the parameter gradients of both networks are given by:

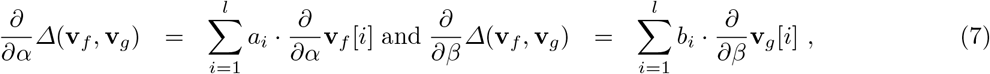

where the gradients of network outputs are obtained via back-propagation and their respective constants are given by *a*_*i*_ = 2 · (1 − **v**_*g*_[*i*]) · (**v**_*f*_ [*i*] − **v**_*g*_[*i*]) + 2*λ* · **v**_*g*_[*i*] · (**v**_*f*_ [*i*] − 1) and *b*_*i*_ = 2 · (**v**_*g*_[*i*] − 1) · (**v**_*f*_ [*i*] − **v**_*g*_[*i*]) − (**v**_*f*_ [*i*] − **v**_*g*_[*i*])^2^ + *λ* · **v**_*g*_[*i*] · (**v**_*f*_ [*i*] − 1).

## 4 Results

### Implementation details

We implement our method using PyTorch and deploy all experiments on a RTX-2060 GPU. Due to limited GPU memory, each training epoch only computes a batch divergence which averages over *N* = 10 randomly sampled subsequences of length *l* = 500 × (*w* + *k*). We set *λ* = 1 and use architectures of PriorityNet and TemplateNet as given in Fig. 2 and Section 3.4 respectively. Network weights are optimized using the ADAM optimizer [9] with learning rate *η* = 5*e*^−3^. Our implementation is available at https://github.com/Kingsford-Group/deepminimizer.

### Comparison baselines

We compare DeepMinimizer with the following benchmarks: (a) random minimizer baseline; (b) Miniception [19]; (c) PASHA [4]; and (d) PolarSet Minimizer [20]. Among these methods, (d) is a sequence-specific minimizer scheme. For each method, we measure the density factor 𝒟 obtained on different segments of the human reference genome: (a) chromosome 1 (Chr1); (b) chromosome X (ChrX); (c) the centromere region of chromosome X [13] (which we denote by ChrXC); and (d) the full genome (Hg38). We used lexicographic ordering for PASHA as suggested by [19]. Random ordering is used to rank *k*-mers within the UHS for Miniception, and outside the layered sets for PolarSet.

### Visualizing the mechanism of DeepMinimizer

First, we show the transformation of the priority scores assigned by ScoreNet and TemplateNet over 600 training epochs. Fig. 3 plots the outputs of these networks evaluated on positions 500 to 1000 of ChrXC, and their corresponding locations of sampled *k*-mers. Initially, the PriorityNet assignment resembles that of a random minimizer and expectedly yields 𝒟 = 2.05. After training, the final TemplateNet assignment converges with a different phase shift than its initial assignment, but its period remains the same. Simultaneously, the PriorityNet assignment learns to match this template, hence induces a visibly sparser sketch with 𝒟 = 1.39. This result clearly demonstrates the negotiating behaviour of our twin architecture to find optimal neighborhood of score assignments.

**Fig. 3.**
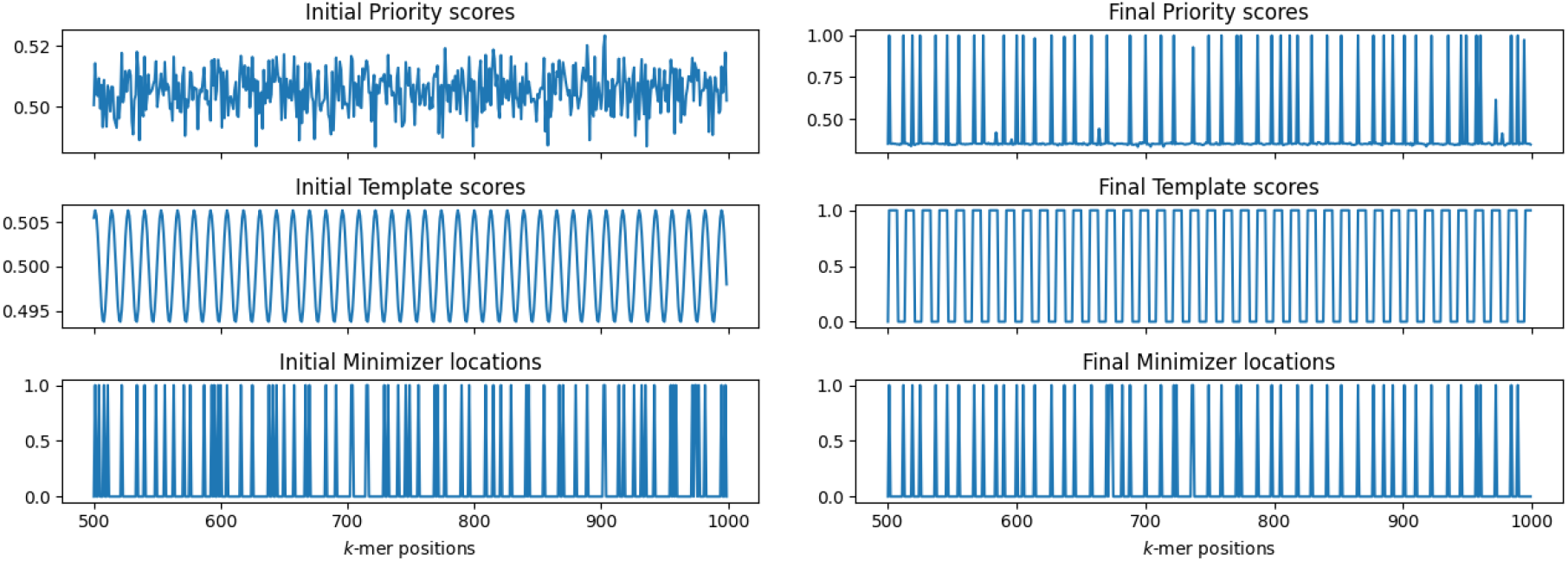
Visualization of PriorityNet and TemplateNet score assignments on positions 500 − 1000 of ChrXC with *w* = 13, *k* = 8. Left: Initial assignments (𝒟 = 2.05); Right: Final assignments after 600 training epochs (𝒟 = 1.39). The bottom plots show corresponding locations of sampled *k*-mers: a value of 1 means selected, and 0 otherwise.

### Convergence of our proxy objective

We further demonstrate that our proxy objective meaningfully improves minimizer performance as it is optimized. The first two columns of Fig. 4 show the best density factors achieved by our method over 600 epochs on two scenarios: (a) varying *k* with fixed *w*; and (b) varying *w* with fixed *k*. The experiment is repeated on ChrXC and Hg38. In every scenario, DeepMinimizer starts with *𝒟 ≃* 2.0, which is only comparable to a random minimizer. We observe steady decrease of *𝒟* over the first 300 epochs before reaching convergence, where total reduction ranges from 11 − 23%.

**Fig. 4.**
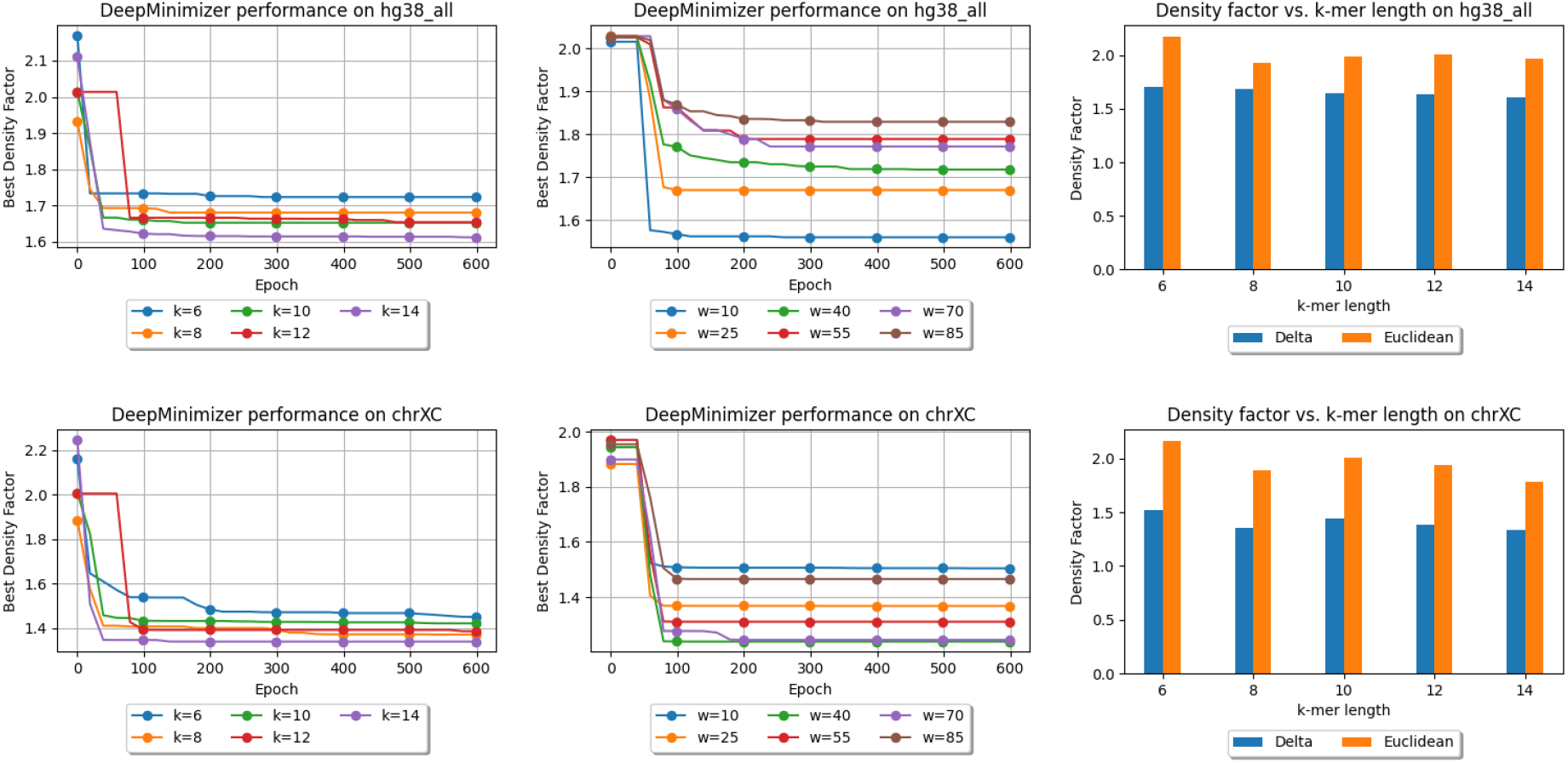
Best density factors obtained by DeepMinimizer on Hg38 (above) and ChrXC (below) over 600 training epochs. Left: fix *w* = 13, and vary *k* ∈ {6, 8, 10, 12, 14}; Center: fix *k* = 14, and vary *w* ∈ {10, 25, 40, 55, 70, 85}; Right: Comparing proposed *Δ*-divergence and *𝓁*^2^-divergence.

Generally, larger *k* values lead to better performance improvement at convergence. This is expected since longer *k*-mers are more likely to occur uniquely in the target sequence, which makes it easier for a minimizer to achieve sparse sampling. In fact, previous results have shown that when *k* is much smaller than log *w*, no minimizer will be able to achieve the theoretical lower-bound 𝒟 [19]. On the other hand, larger *w* values lead to smaller improvements and generally slower convergence. This is because our ensemble parameterization of TemplateNet scales with the window size *w* and becomes more complicated to optimize as *w* increases.

### Evaluating our proposed divergence measure

The last column of Fig. 4 shows the density factors achieved by our DeepMinimizer method, respectively specified by the proposed divergence function in Eq. 6 and *𝓁*^2^-divergence. Here, we fix *w* = 14 and vary *k* ∈ {6, 8, 10, 12, 14} and observe that with the *𝓁*^2^-divergence, we only obtain performance similar to a random minimizer. On the other hand, with our divergence function, DeepMinimizer obtains much lower densities on all settings, thus confirming the intuition in Section 3.5.

### Comparing against other minimizer selection benchmarks

We show the performance of Deep-Minimizer compared to other benchmark methods. DeepMinimizer is trained for 600 epochs to ensure convergence, as shown above. Fig. 5 shows the final density factors achieved by all methods, again on two comparison scenarios: (a) fix *w* = 13, and vary *k* ∈ {6, 8, 10, 12, 14}; and (b) fix *k* = 14, and vary *w* ∈ {10, 25, 40, 55, 70, 85}.

**Fig. 5.**
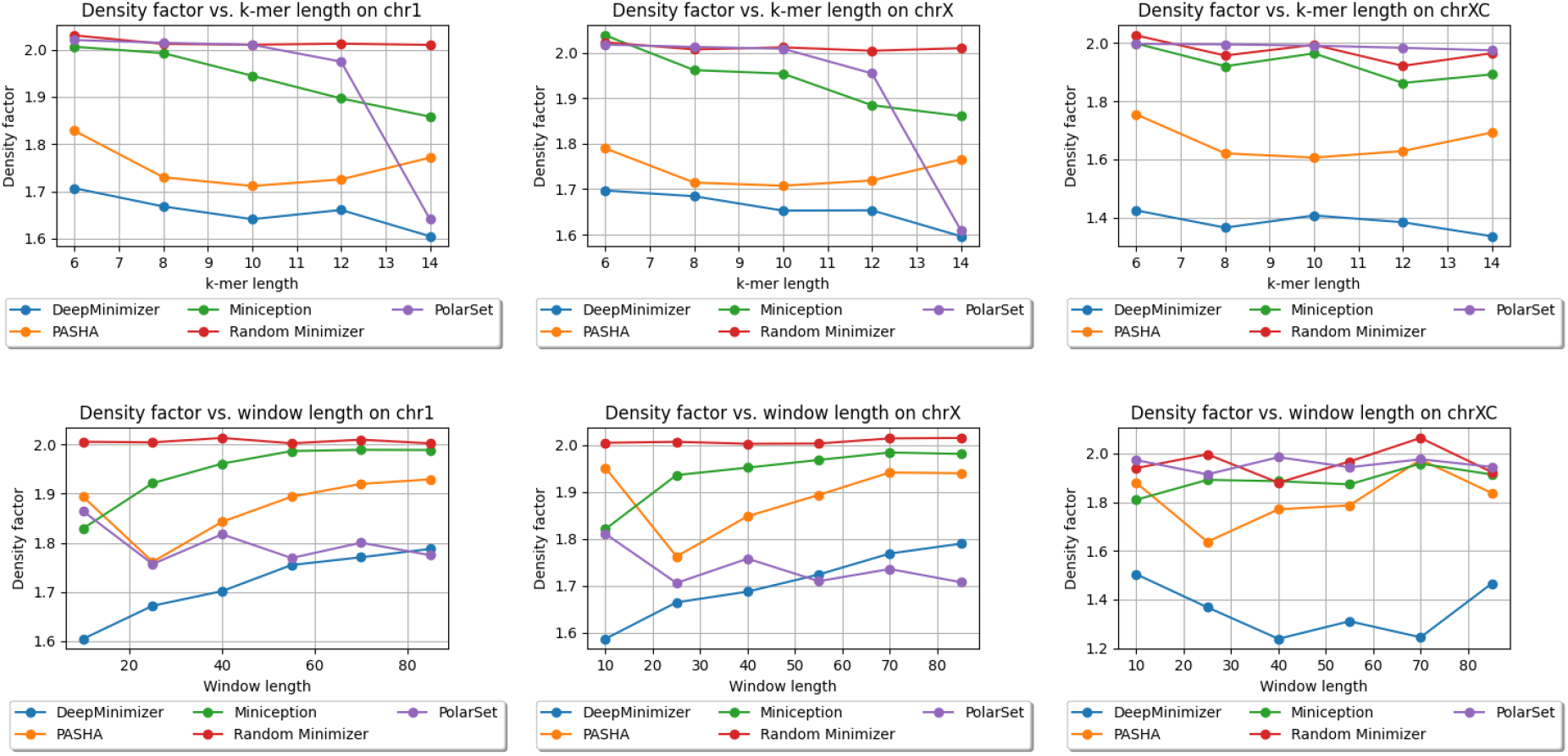
Density factors obtained by DeepMinimizer (600 training epochs), Random Minimizer, PASHA, Miniception and PolarSet on Chr1, ChrX and ChrXC. Above: fix *w* = 13, and vary *k* ∈ {6, 8, 10, 12, 14}; Below: fix *k* = 14, and vary *w* ∈ {10, 25, 40, 55, 70, 85}.

DeepMinimizer consistently achieves better performance compared to *non-sequence-specific* minimizers (i.e., PASHA, Miniception) on all settings. We observe up to 40% reduction of density factor (e.g., on ChrXC, *w* = 70, *k* = 14), which clearly demonstrates the ability of DeepMinimizer to exploit *sequence-specific* information. Furthermore, we also observe that DeepMinimizer outperforms our *sequence-specific* competitor, PolarSet, in a majority of settings. The improvements over PolarSet are especially pronounced for smaller *k* values, which are known harder tasks for minimizers [19]. On larger *w* values, our method performs slightly worse than PolarSet in some settings. This is likely due to the added complexity of optimizing TemplateNet, as described in convergence ablation study of our method.

In addition, we also conduct investigation on the centromere region of chromosome X (i.e., ChrXC), which contains highly repetitive subsequences [6] and has been shown to hamper performance of PolarSet [20]. Fig. 5 shows that PolarSet and the UHS-based methods perform similarly to a random minimizer, whereas our method is consistently better. Moreover, we observe that DeepMinimizer obtains near-optimal densities with ChrXC on several settings. For example, we achieved 𝒟 = 1.22 when *k* = 14, *w* ∈ {40, 70}, which is significantly better than the results on Chr1 and ChrX. This suggests that ChrXC is not necessarily more difficult to sketch, but rather good sketches have been excluded by the UHS and polar set reparameterizations, which is not the case with our framework.

### Runtime performance

DeepMinimizer runs efficiently with GPU computing. In all of our experiments, each training epoch takes approximately 30 seconds to 2 minutes, depending on the choice of *k* and *w*, which controls the batch size. Performance evaluation takes between several minutes (ChrXC) to 1 hour (Hg38), depending on the length of the target sequence. Generally, our method is cost-efficient without frequent evaluations. Our most cost-intensive experiment (i.e., convergence ablation study on Hg38) requires a full-sequence evaluation every 20 epochs over 600 epochs, thus takes approximately 2 days to complete. This is faster than PolarSet, which has a theoretical runtime of 𝒪 (*n*^2^) and takes several days to run with Hg38. A more detailed runtime ablation study on Chr1 is provided in Appendix B.

## 5 Conclusion

We introduce a novel framework called DeepMinimizer for learning *sequence-specific* minimizers. This is achieved via casting minimizer selection as optimizing a *k*-mer scoring function *ρ*. We propose a more well-behaved search space for minimizers, given by a neural network parameterization of *ρ*, called PriorityNet. Then, we introduce a complementary network, called TemplateNet which pinpoints optimal scoring templates and guides PriorityNet to the neighborhood of low-density assignments around them. Coupling these networks leads to a fully differentiable proxy objective that can effectively leverage gradient-based learning techniques. DeepMinimizer obtains better performance than state-of-the-art sequence-agnostic and sequence-aware minimizer selection schemes, especially on known hard tasks such as sketching the repetitive centromere region of Chromosome X. However, we also observe mild limitations in several settings with large window length *w*, which hampers the performance of DeepMinimizer. This is likely due to the heuristic construction of our TemplateNet component, which we will investigate in our future work.

## Acknowledgements

This work was supported in part by the Gordon and Betty Moore Foundation’s Data-Driven Discovery Initiative [GBMF4554 to C.K.], by the US National Institutes of Health [R01GM122935], and the US National Science Foundation [DBI-1937540]. Conflict of Interest: C.K. is a co-founder of Ocean Genomics, Inc.

## A Proof of Proposition 1

We first re-express the density factor of *S* in terms of a priority score assignment **v** ∈ [0, 1]^*l*^. Note that this expression will hold regardless of whether **v** satisfies the consistency constraint (Section 3.2). Particularly, let *γ*_1_ = 1 and 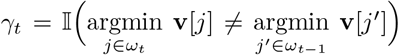 indicates the event the *t*-th window picks a different *k*-mer than the (*t* − 1)-th window, we have:

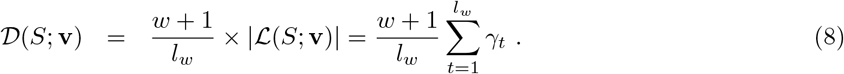

Without loss of generality, we assume 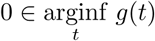 since this can always be achieved via adding a constant phase shift to *g*. As *g* has a fundamental period of *w*, this implies {*uw* | *u* ∈ ℕ} *⊆* arginf_*t*_ *g*(*t*), which further reduces to {*uw* | *k* ∈ ℕ} = arginf_*t*_ *g*(*t*) when condition (2) holds.

Let us now derive the values of *γ*_*t*_ for *t* ∈ *ℐ*_*u*_ ≜ [(*u* − 1)*w* + 1, *uw*], *u* ∈ ℕ^+^. We have:

– *uw* ∈ arginf *g*(*t*),
– *∀t* ∈ *ℐ*_*u*_ such that *t ≠ uw*, we have 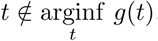, and
– *∀t* ∈ *ℐ* (*u*) : *uw* ∈ *ω*_*t*_, which follows from the above argument and the definition of window *ω*_*t*_.

Together, these observations imply that 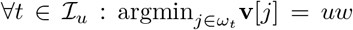 and consequently *γ*_*t*_ = 0 for all values of *t* ∈ *ℐ*_*u*_ except *t* = (*u* − 1)*w* + 1. For *u* = 1, we trivially have *γ*_(*u*−1)*w*+1_ = 1 by definition of *γ*_1_. For *u >* 1, we have 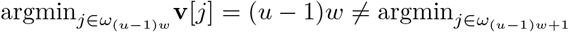, which also implies *γ*_(*u*−1)*w*+1_ = 1. Following the above derivations, we have:

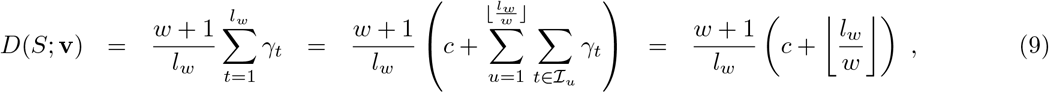

where the third equality follows from the derived values *γ*_*t*_ for *t* ∈ *ℐ*_*u*_. Finally, using the fact that *c* = 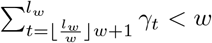 and the sufficient length assumption *l*_*w*_ *≫ w*^2^, we have:

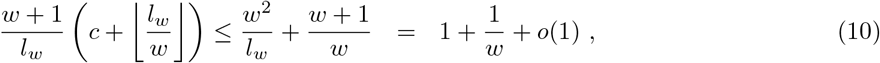

which concludes our proof. □

## B Other Empirical Results

This section contains extra experiments that showcase various aspects of our DeepMinimizer framework. For all experiments, we use the same implementation, benchmarks and settings as detailed in Section 4.

### Density performance of DeepMinimizer on more sequence baselines

We deploy DeepMinimizer on Chr1 and ChrX. For both sequences, we observe the best density factor obtained over 600 training epochs for various values of *k* and *w*. Fig. 6 shows that DeepMinimizer consistently improves density factors until convergence, which tends to happen between 200 − 300 training epochs for all experiments.

**Fig. 6.**
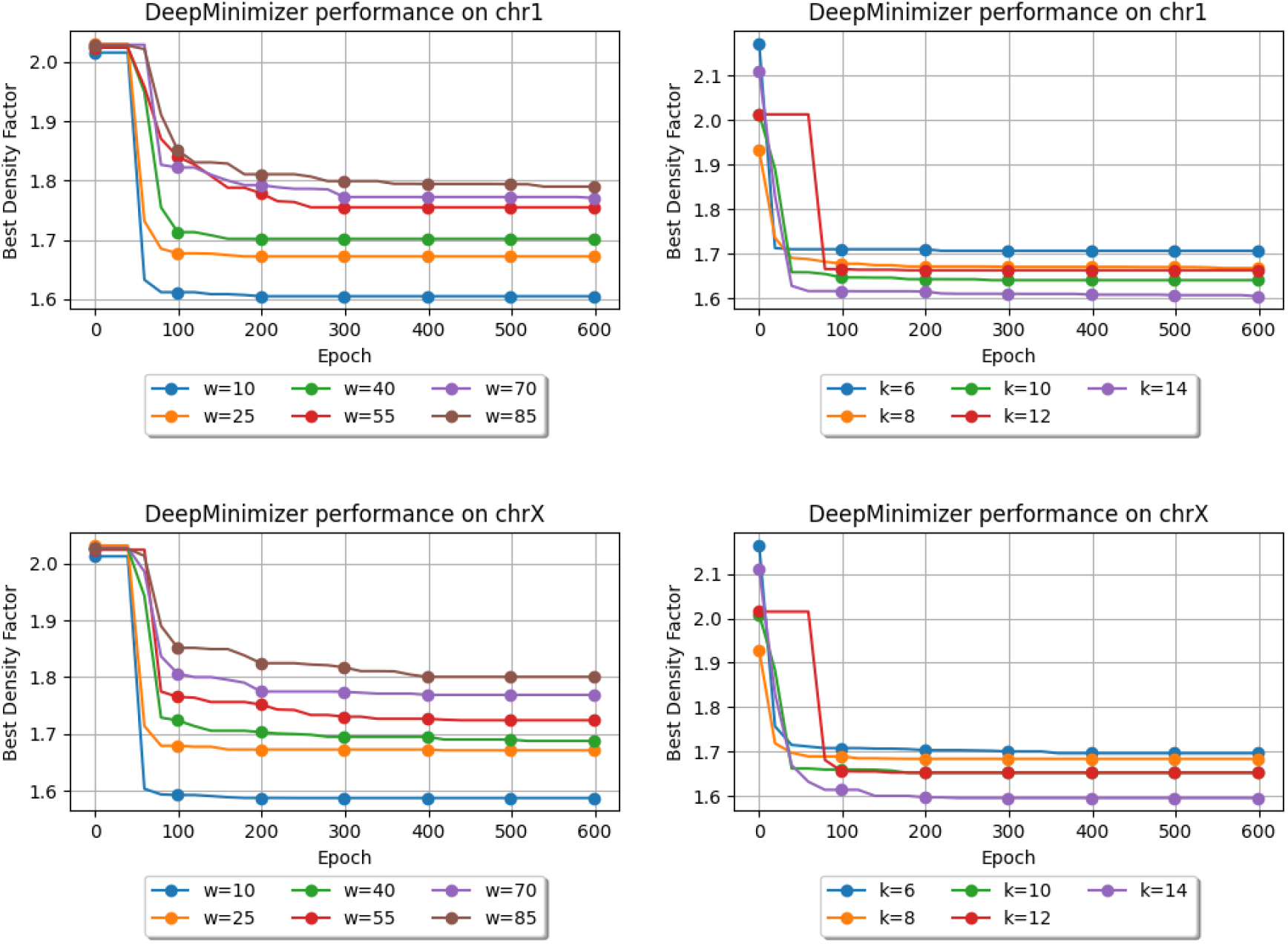
Demonstrating convergence of DeepMinimizer on Chr1 (left) and ChrX (right) with different *w, k* values.

### DeepMinimizer outperforms other baselines on large sequences

Fig. 7 compares the performance of DeepMinimizer and various comparison baselines on the entire human genome Hg38. We measure the best density factor obtained over 600 training epochs for various values of *k* and *w* and observe that DeepMinimizer consistently achieves the best performance among comparison baselines.

**Fig. 7.**
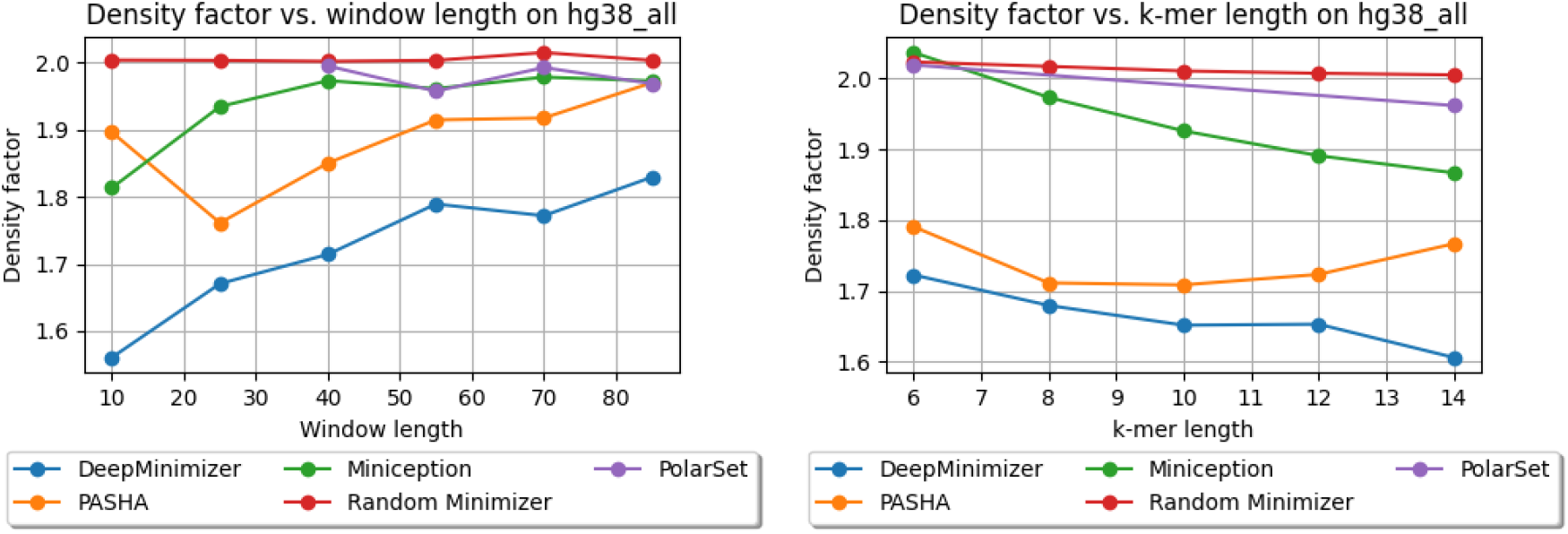
Comparing performance of DeepMinimizer with other benchmarks on Hg38 for different values of *w, k*.

### Density performance of DeepMinimizer on large values of *k*

Fig. 8 (left) showcases the performance of DeepMinimizer on Chr1 with large values of *k*. We fix *w* = 13 and observe the best density factor obtained over 600 training epochs for various values of *k* up to 320. We show that DeepMinimizer behaves similarly for large *k*, and achieves the best density *𝒟* = 1.22 with *k* = 160.

**Fig. 8.**
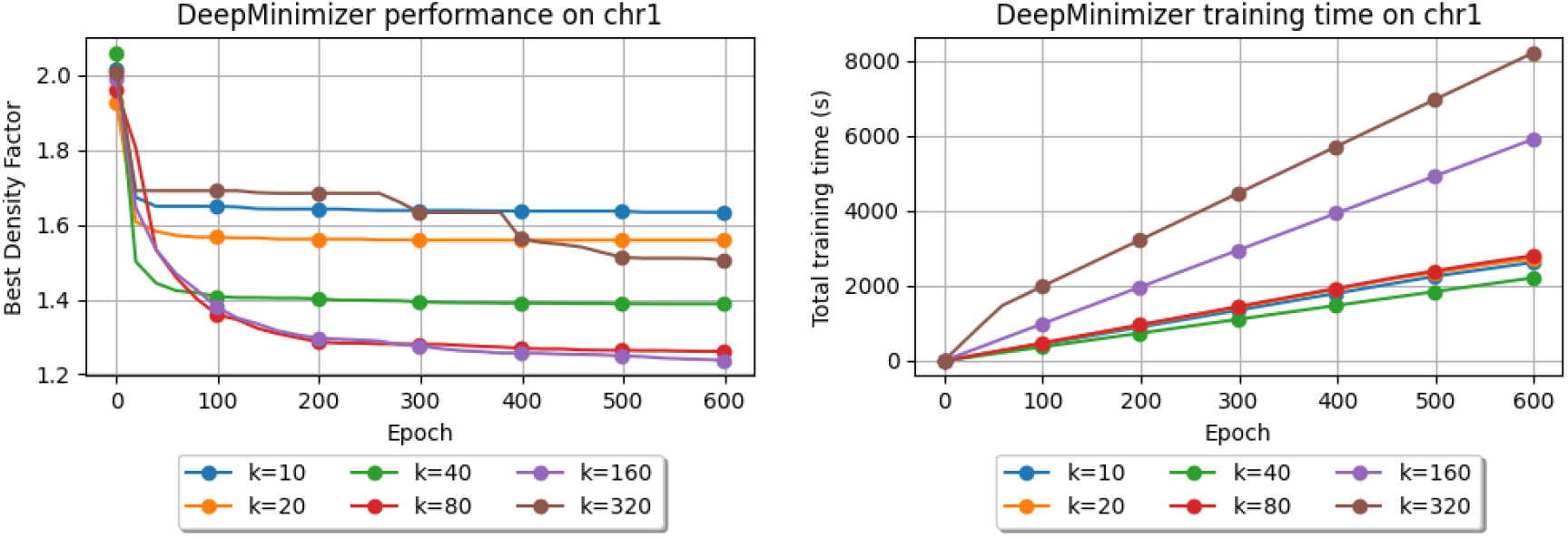
Best density obtained (left) and runtime (right) of DeepMinimizer for *k* ∈ {10, 20, 40, 80, 160, 320} on Chr1.

### Runtime performance of DeepMinimizer on large values of *k*

Fig. 8 (right) measures runtime (in seconds) of DeepMinimizer on Chr1 over 600 epochs. Larger *k* values require PriorityNet to have more parameters. Expectedly, we observe runtime for *k* = 40, 80, 160, 320 to increase in the same order. For *k* = 10 and 20, however, the runtimes are approximately the same as *k* = 80. We note that a smaller *k* value means there are more *k*-mers in the same sequence. As such, even though PriorityNet is more compact for these values of *k*, we will incur some overhead from querying it more often.

## Notes

### Competing Interest Statement

The authors have declared no competing interest.

